# 3D Microscale Mechanical Simulations of Hydrogel Coated Electrospun Meshes

**DOI:** 10.64898/2026.01.19.700377

**Authors:** Evan He, Shruti Motiwale, Elizabeth Cosgriff-Hernandez, Michael S. Sacks

## Abstract

Electrospun fiber meshes have long served as biomaterials in a wide range of biomedical applications due to their functional similarities to extracellular matrix and highly tunable properties. Altering the mechanical behaviors of individual fibers and their microarchitecture (e.g.; diameter, crimp, orientation, density) can in principle be used to control bulk level behaviors. Moreover, electrospun meshes are often combined with softer coatings and hydrogels to control surface interactions with body tissues. Yet, fully optimizing their behaviors for specific applications remains an elusive target due to a continued lack of understanding of the micromechanical mechanisms and their relation to bulk mechanical behaviors. Our goal herein was to understand how actual nanoCT-generated 3D microfiber geometry can be used to predict bulk mechanical properties of hydrogel-mesh composites. Electrospun polyurethane meshes were fabricated with a random fiber orientation and coated with a PEG-based hydrogel. The fiber-hydrogel composite was then imaged with a nanoCT scanner at a voxel resolution of 180 nm. From these images, custom Python programs were written to segment, refine, and tesselate a high-resolution finite element of the fiber mesh and hydrogel volumes into a single integrated bi-material finite element model. The resulting mesh was used to run simulations of the planar biaxial mechanical tests used to characterize the bulk mechanical behaviors. Our framework thus enabled systematic investigations of both the macroscopic bulk mechanical response of the overall fiber mesh and the microscopic localized mechanical response of fibers under various stages of loading. The resultant simulations were accurate and predictive of the bulk mechanical responses. It is interesting to note that the fiber-hydrogel composite material experienced the largest stresses within the fiber phase and the largest strains within the hydrogel. This key result underscores that while the previous analytical model assumed local affine deformations, at the microscale this assumption does not hold. We also found very different effective fiber stress-strain responses in each model. It is likely these differences are due to the substantial heterogeneous non-affine local deformations present in the actual fiber-hydrogel composite. This finding further reveals the need for more rigorous approaches to better understand how electrospun-based materials function in order to improve their use in modern medical devices and implants.

## 1. INTRODUCTION

Electrospun fiber meshes have long served as functional biomaterials in a wide range of biomedical applications, due to their functional similarities to natural extracellular matrices (ECM) and highly tunable properties. The tunability of electrospun mesh mechanical behaviors is a direct result of the flexibility of their fabrication methods. Electrospun meshes are generated by applying electrostatic forces to polymer solutions, drawing fibers with diameters ranging from the nano-to microscale. The resulting structures are highly porous fibrous meshes, generally comprising approximately 30% fiber volume fraction. The final mesh architecture—including fiber diameter, orientation, alignment, and density—is sensitive to multiple processing parameters, including solution rheology, applied voltage, collector geometry, and ambient humidity [1]. Bulk mesh properties can be readily tuned by tailoring the mechanical behavior of individual fibers, as well as by controlling fiber microarchitecture [2, 3].

Given the large number of fabrication parameters and the complexity of the resulting mechanical behaviors, there has been significant efforts in developing electrospun mesh structure-mechanical models. For example, Courtney et al. [4] evaluated electrospun polyurethane meshes using a structural-based constitutive model derived from similar soft tissue models that incorporated fiber orientation. This early approach revealed quantitatively strong reliance on bulk fiber orientation and the resulting anisotropic non-linear mechanical behaviors. Comparisons with experimental data were also conducted and revealed close agreement between model derived and measured fiber orientations. Sylianopoulos et al. [5] developed a multiscale approach to link fiber-level structure (fiber diameter, fiber orientation, and fiber density) to bulk mechanical response. More aligned fiber meshes were found to produce greater anisotropy, while less aligned fiber meshes exhibited lower, more uniform mechanical responses. D’Amore et al. [6] developed an image-based approach to collect fiber angle distribution, fiber connectivity, fiber intersection density, and fiber diameter to characterize fiber network topology from SEM images. Increased mandrel velocity during electrospinning was found to produce more aligned fibers fewer fiber intersections, and larger fiber diameters. Carleton et al. [7] developed 2D statistical fibrous network-based models that utilized stochastic fiber density, fiber intersection density, and orientation distribution functions [8]. They were able to perform parametric simulations that captured the effects of fiber tortuosity, alignment, as well as describe local non-affine behaviors. Rizvi et al. [9] developed a model using a probability density function to describe how the microstructural properties of a fibrous matrix (e.g. fiber orientation, stiffness, diameter, curvature, alignment) contributed to the bulk mechanical mesh properties.

Collectively, these works demonstrated that electrospun meshes can be modeled and their responses directly related to bulk structural measures. However, most extant experimental data exclusively utilized extensional loading paths in either uniaxial or planar biaxial modes, leaving out in-plane shear modes. However, in many applications electrospun meshes are subjected to in-plane shearing deformations (e.g. [10]). Based on these findings, Motiwale et al. [11] performed planar biaxial mechanical evaluations on hydrogel-coated polyurethane fiber meshes that included both extensional and in-plane shear measurements. An extended meso-structural constitutive model was developed to relate key structural features (fiber orientation) to the resultant generalized in-plane deformations [12]. For extensional deformations a constitutive model form similar to Courtney et al. [4] was found to be quite sufficient, yet was unable to account for shear deformation responses. A fiber-fiber interaction term, based on related work on cross-linked pericardium [13], was then developed. The extended model form was found to be in excellent agreement with experimental results for all biaxial deformation modes. This work suggested the presence of significant fiber-fiber interactions in electrospun meshes mechanical responses.

Regardless of the specifics, all the above electrospun mesh constitutive models can predict the macroscopic mechanical response of electrospun fibers reasonably well. Yet it remains to be seen how the actual microscale 3D fiber geometry dictates macro-scale behaviors. Such detailed 3D models will be needed to better design electrospun meshes for specific applications, as well as determine mechanisms of damage and fatigue. Further, we and others have also utilized hydrogel coated electrospun meshes, which might induce additional fiber-gel interactions. The evidence for potential gel-fiber and fiber-fiber interactions should thus also be evaluated to better understand their contributions to the overall mechanical response. Although the work by Carleton [7] included fiber-fiber bonding, this work was limited to two-dimensional (2D) simulations only, as three-dimensional (3D) random walk methods remain difficult to utilize and quantitative 3D geometry data is currently lacking. Taken as a whole, present approaches remain limited in their lack of the ability to account for full 3D micro-mechanical mechanisms.

In the present work, we developed a full 3D micromechanical model of electrospun meshes under biaxial loading to directly generate bulk mechanical behaviors. Our goal was to understand how actual 3D fiber geometry at the microscale level impacts the bulk mechanical properties of fibrous electrospun meshes. We first developed a pipeline to transform a 3D nanoCT reconstruction of an electrospun fibrous mesh specimen into a high fidelity 3D finite element model that captured the structure of all individual fibers. This framework enabled systematic investigation of both the macroscopic bulk mechanical response of the overall fiber mesh and the microscopic localized mechanical response of fibers under various stages of loading. The resultant simulations were then compared to experimental data by [12] and novel insights gleaned from the modeling results.

## 2. METHODS

### 2.1. Summary of Approach

The pipeline began with nanoCT imaging of an electrospun mesh. The resulting images were then binarized to extract the fibers. To generate finite element meshes from these images, a marching cubes and Delaunay algorithms were then applied in a two-step process to create surface and volume meshes, respectively, of varying sizes. The resulting volume meshes were then imported ABAQUS/Standard finite element solver, and simulations performed to emulate the experimental n-plane deformation modes. The resulting simulated effective bulk material behaviors were then calibrated to the experimental data [11]. (Figure 1)

**Figure 1.**
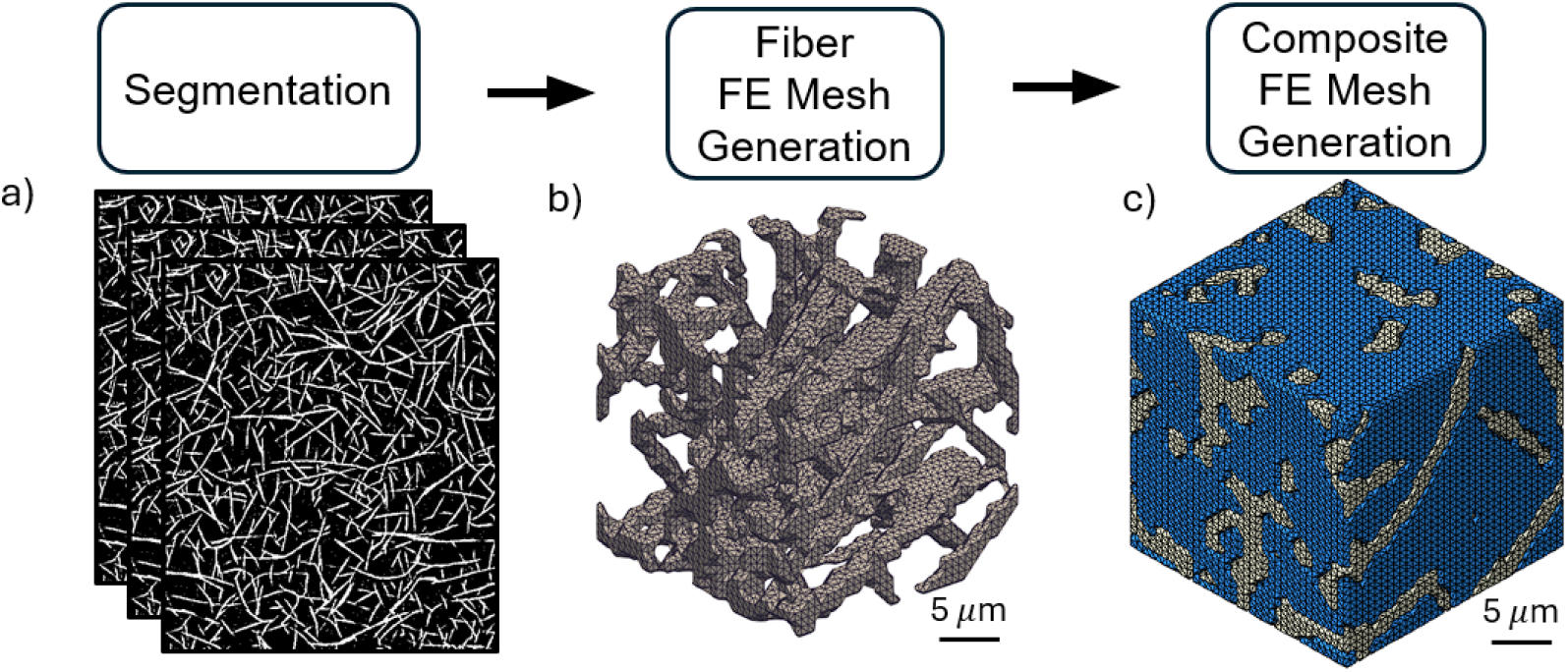
The overall project image-to-model pipeline. a) ImageJ is used to process nanoCT image data. b) A two-step application of the marching cubes algorithm and Delaunay triangulation algorithm are applied to the processed images to create finite element meshes of the electrospun fiber geometry and the negative space around it. c) The two meshes are stitched together to represent a fiber-hydrogel composite mesh.

### 2.2. Electrospun mesh fabrication

All experimental mechanical data of electrospun mesh and fiber-hydrogel composite were taken from Motiwale et al. [12]. Briefly, the electrospun polyurethane mesh was fabricated using a 25 wt% solution of Bionate® 80A in dimethylacetamide. The solution was loaded into a syringe with a 20G needle and dispensed using a syringe pump at a constant flow rate of 0.5 mL/h. A positive voltage of 15 kV was applied to the needle and electrospun fibers were collected on a negatively-charged mandrel (−5 kV, 4 cm diameter, 3.5 cm length, 50 cm collection distance) rotating at 50 RPM. Electrospinning was performed at ambient conditions (22–25 C, 45%–55% relative humidity). For the hydrogel coatings, precursor solutions were first prepared by dissolving polyethylene glycol diacrylate (PEGDA, 3.4, 6, or 10 kDa) at 10 wt% (w/vol) in deionized water with Irgacure 2959 (0.1 wt%). The bulk hydrogel was prepared by pipetting the solution between 1.5 mm spaced plates and UV crosslinking (365 nm, 4 mW/cm2). The fiber-hydrogel composite was prepared by adding the electrospun fiber mesh (12×7 mm) between 500 µm spacer plates after a graded ethanol/water soak and UV crosslinking for 6 min on both sides. Prior to testing, bulk hydrogels and composites were immersed in deionized water for 24 h to allow for equilibrium swelling prior to characterization.

### 2.3. Nano-CT Imaging and Processing

The electrospun fiber mesh was imaged with a General Electric v|tome|x L300 multi-scale nano/microCT scanner (Center for Quantitative Imaging, Pennsylvania State University) with a voxel resolution of 180nm. The resulting nanoCT image stack contained 1628 images; and each image had pixel dimensions of 1919 x 2006, resulting in a final imaging dimension of 346.9 *µm* x 362.7 *µm* x 347.6 *µm*. The cuboidal dimensions of the specimen were about 293 *µm* x 232 *µm* x 216 *µm* (since the specimen did not fill the entire imaging view) (Figure 2). Using ImageJ, individual fiber structure was isolated by binarizing the nanoCT images using a grayscale threshold set to 170 [14]. This threshold value was chosen because it best retained visible fiber morphology while minimizing loss of overall fiber structure. Since there was also a misalignment between the imaging axis and the specimen axis, a rotation was applied to all images to best align the fiber plane with the imaging plane. Additionally, the images were denoised using the “Erode” function to remove unassociated lone pixels (background noise). The resulting images represented the electrospun fiber mesh structure as white pixels and the empty background space as black pixels (Figure 1a).

**Figure 2.**
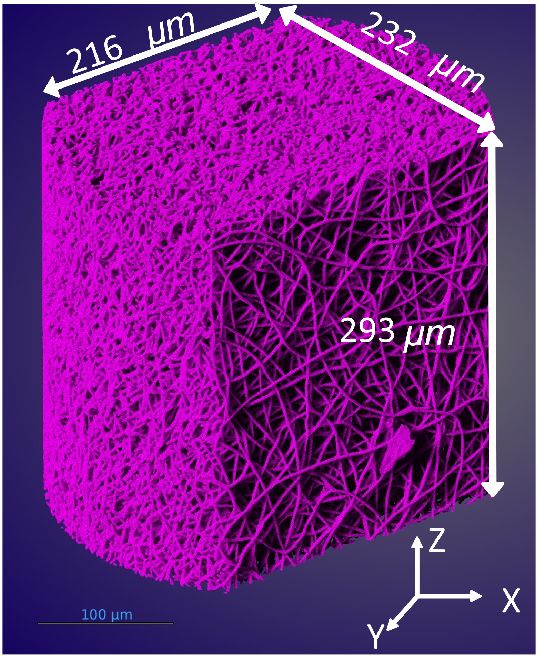
3D rendering of an ES mesh sample reconstructed from nanoCT images, which was 293 *µm* x 232 *µm* x 216 *µm*

### 2.4. RVE Analyses

An essential part of our study was determination of the representative volume element (RVE) size to insure the present micro model properly represented the ES microstructure. This was done using a convergence analysis for fiber volume fraction as follows. Formally, an RVE is the smallest unit necessary to represent a heterogeneous material [15], [16]. A proper RVE size thus ensures that the effective behavior of a material volume is representative of the material as a whole [17]. To determine the minimal RVE size to limit computational cost, different candidate sizes were evaluated [18]. RVE size was determined by identifying the candidate size at which the computed fiber volume fractions converged. The convergence analysis was performed using incrementally larger cubic volume subsets up to the largest possible cubic volume of 200 *µm*^3^ (Figure 3). The fiber volume fraction was calculated by dividing the number of pixels considered a fiber with the total number of pixels in the sub-cube’s volume.

**Figure 3.**
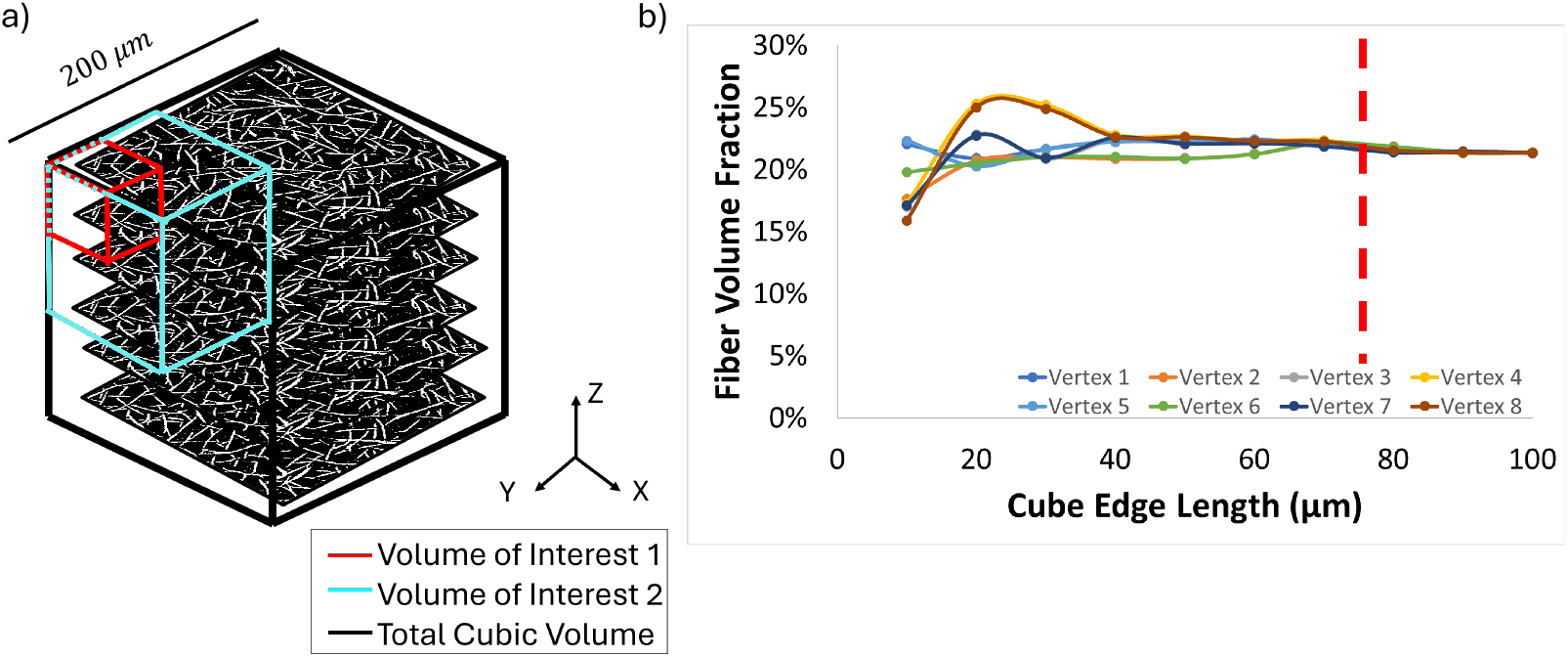
RVE convergence study. a) A total cube size of 200 *µm* x 200 *µm* x 200 *µm* is used to create sub-cubes (volumes of interest) from each of the total cube’s eight vertices. b) Fiber volume fractions are plot for each volume of interest. Convergence of fiber volume fractions was found around a 75 *µm*^3^ volume of interest.

Fiber volume fraction of the entire binarized nanoCT image dataset was found to be 22.14%. Fiber volume fractions for the different-sized cubic volume subsets converged around a cube size of 75 *µm*^3^, with an average fiber volume fraction of 22.10% *±*0.001%. The 25 *µm*^3^ cubic mesh had an average fiber volume fraction of 22.27% *±*0.018%, and the 50 *µm*^3^ cubic mesh had an average fiber volume fraction of 21.95% *±*0.006%. This analysis confirmed that the 75 *µm*^3^ image volume was sufficiently large to capture the micro-structural characteristics of the full nanoCT dataset. Following these results, the 75 *µm*^3^ cubic mesh was chosen as the optimal RVE size for the remainder of the study (Figure 3).

### 2.5. Finite element model

#### Mesh generation

The Dragonfly 3D World program (Dragonfly, Montreal, Quebec) was used to generate a surface mesh from the nanoCT image stack with triangle elements using the marching cubes algorithm [19]. First, the nanoCT image stack was imported into Dragonfly. To create different cube sizes, the nanoCT image stack was cropped concentrically from the center of the volume to avoid edge artifacts, resulting in 3 different cube sizes: 25 *µm*^3^, 50 *µm*^3^, and 75 *µm*^3^. Two regions of interest were defined by using a watershed transform to threshold the cropped image stacks. A watershed transform segments an image using pixel intensity values by treating them as a topographic surface, where local minima correspond to catchment basins, and ridges define the boundaries between neighboring regions [20]. The voxels in the first region of interest represented the electrospun fibers in the nanoCT images, while the second region of interest represented the empty space surrounding the fibers. For this study, the first region of interest was processed as a volume to represent electrospun fibers, while the second region of interest was processed as a volume to represent a hydrogel material; both were later combined together into a fiber-hydrogel composite. Using the marching cubes algorithm, a surface mesh containing triangle elements was generated for each region of interest. Thus, for cubic volumes of each size, there were 2 associated surface meshes that were exported for further processing: a fiber surface mesh and a hydrogel surface mesh.

The output surface mesh from Dragonfly was then imported into GMSH to generate a volume mesh with linear tetrahedral elements using the Delaunay algorithm [21]. First, the fiber surface mesh was imported into GMSH, then a physical surface group was defined for the fiber mesh. The physical surface group contained all the surface elements grouped together [21]. Next, a surface loop and volume definition was applied to the fiber mesh. A surface loop defines the exterior boundary for a volume, and a volume definition initializes the three-dimensional space required for volume meshing [21]. Then, the gel surface surface mesh was imported alongside the fiber surface mesh using the “Merge” function in GMSH. A surface loop and volume definition was also applied separately to the hydrogel surface mesh. Both surface meshes were volume meshed simultaneously by applying the “3D” mesh function in GMSH to produce linear tetrahedral elements. Local refinement of elements with poor aspect ratios (*>*10) was implemented using the NETGEN module. Finally, the volume meshes were exported as ABAQUS input files.

Since the nanoCT images were defined in the micrometer range, the resulting node coordinates of the meshes and subsequent material model calculations for defining the material model in Abaqus would contain many leading zeros, increasing the risk of numerical precision issues. Thus, to preprocess the cubic meshes, custom Python programs were written to perform the following: 1) scale the Cartesian coordinate system used for node locations, 2) center the mesh to the origin of the coordinate system, and 3) merge the disconnected fiber and hydrogel volume meshes together. The additional step for centering the mesh to the origin of the Cartesian co-ordinate system simplified how boundary conditions were later applied to the mesh. The fiber and hydrogel volume meshes were also combined by merging duplicate shared nodes between all elements to represent a physically bonded mesh (Figure 4).

**Figure 4.**
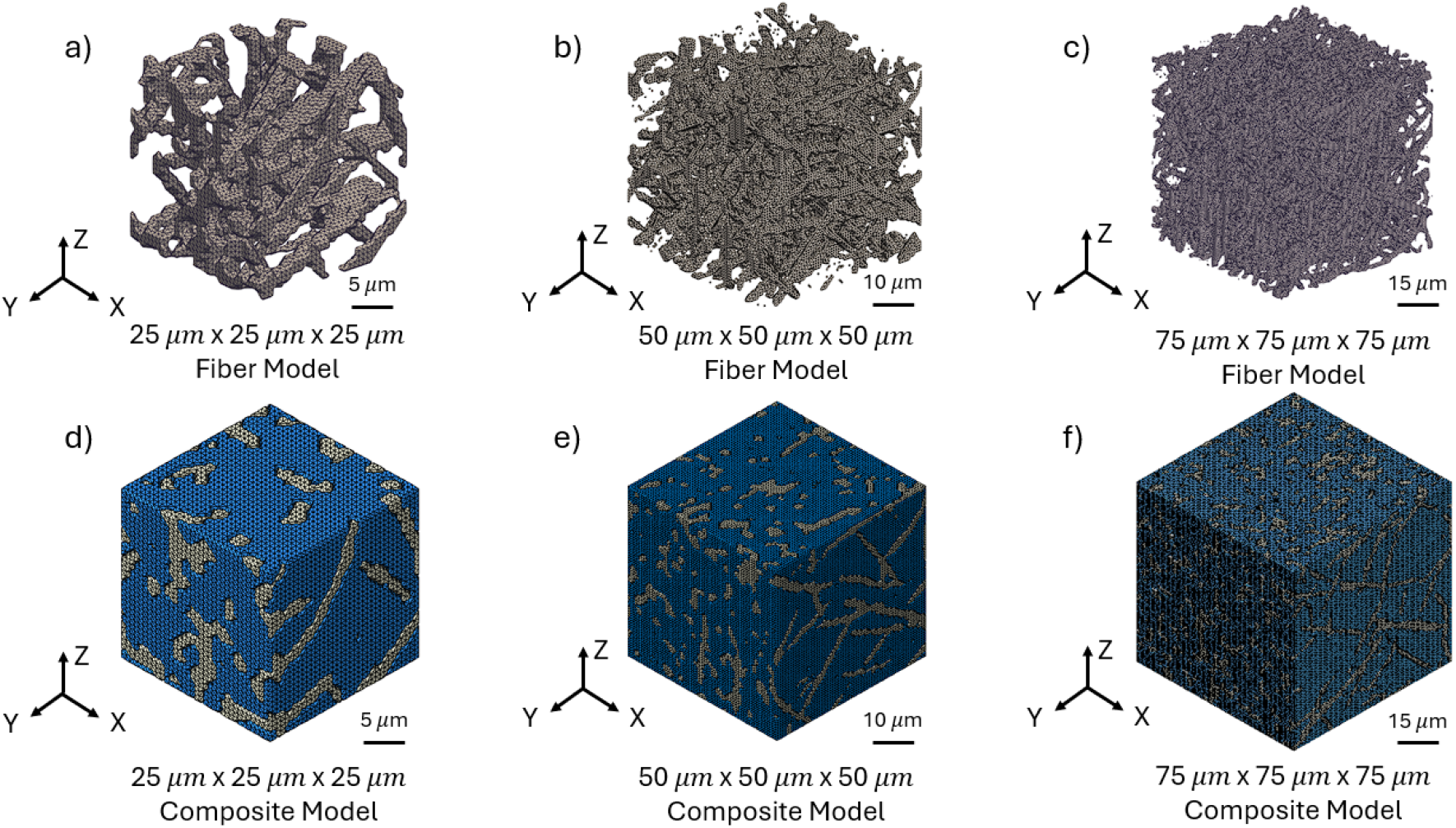
Varying fiber and fiber-hydrogel mesh sizes. a) 25 *µm*^3^ cubic fiber mesh. b) 50 *µm*^3^ cubic fiber mesh. c) 75 *µm*^3^ cubic fiber mesh. d) 25 *µm*^3^ cubic fiber-hydrogel mesh; contains 189,000 elements and 50,000 nodes. e) 50 *µm*^3^ cubic fiber-hydrogel mesh; contains 1.569 million elements and 406,000 nodes. f) 75 *µm*^3^ cubic fiber-hydrogel mesh; contains 4.640 million elements and 1.240 million nodes.

#### Software and boundary conditions

All finite element simulations were performed in ABAQUS/CAE (2023) with the ABAQUS/Standard implicit solver, using, the full Newton-Raphson iterative approach with automatic time increments [22]. All elements of the generated volume meshes were defined as first-order four-node tetrahedral (C3D4) solid elements. Boundary conditions were chosen to replicate the planar biaxial extension tests as performed in [12]. The RVE mesh was oriented such that its faces were normal to the global *x*_1_, *x*_2_, and *x*_3_ coordinate axes. The four faces normal to the *x*_1_ and *x*_2_ directions were selected for displacement loading, and displaced uniformly away from the center of the cubic mesh in their respective direction. The two faces normal to the *x*_3_ direction were traction-free to allow free deformation in this direction. Under these defined boundary conditions, a linear ramping displacement was applied to each of the displaced cubic mesh faces. Three biaxial extension protocols were implemented: 1:1, 1:3, and 3:1. The Python program applied these stretch ratios by scaling the normal and lateral displacement magnitudes according to these modes to achieve a stretch of 1.1:1.1, a stretch of 1.10:1.30, and a stretch of 1.3:1.1. All nodes on each planar face were displaced equally. In pilot studies some initial convergence issues were present due to high aspect ratios (*>*5) for some elements. These elements were identified and the local regions in the vicinity of the offending element re-meshed to avoid further mesh-based issues.

### 2.6. Material Models

#### Hydrogel

The mechanical behavior of the hydrogel was represented using an isotropic Neo-Hookean material model using the following strain energy density function

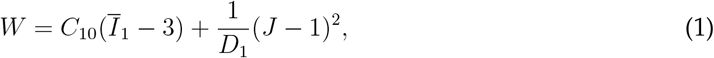

where 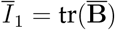 is the first invariant of the deviatoric left Cauchy-Green strain tensor 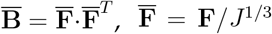 is the deviatoric from of the deformation gradient tensor **F**, and *J* = *det*[**F**]. The model as two material parameters *C*_10_, which is related to the small strain shear modulus, and *D*_1_ which governs compressibility via a penalty term. Values for the hydrogel material parameters were set to *C*_10_ = 42.94 *kPa* and *D*_1_ = 219.3 *kPa*^−1^, taken from [12].

#### Fiber model

In our previous work we utilized the following “analytical” model strain energy Ψ^*sf*^

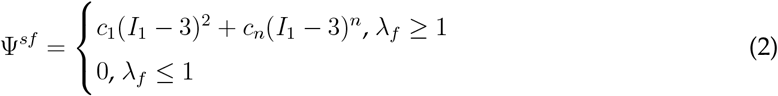

where *c*_1_ = 2079.91 kPa, *c*_*n*_ = 183.31 kPa, and *n* = 8.74 [12]. Since we anticipated possible differences in mechanical responses, for the electrospun fiber phase we utilized standard Yeoh model. The mechanical behavior of the fibers was thus represented using the following general isotropic Yeoh material model

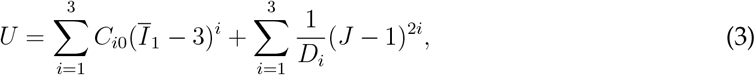

where *C*_*i*0_, *D*_*i*_ are material constants.

### 2.7. Simulations and calibration

#### Planar biaxial simulations

Planar biaxial extension simulations were performed using ratios of axial strains of 1:1, 1:3, and 3:1. To calibrate the fiber model parameters, we first fit Eqn. 3 to simulated stress-strain data using Eqn. 2. We then compared the experimental stress-strain data for the 1:1 biaxial extension loading path to the effective stress-strain results from the simulations. While not very close, this method produced reasonable ‘ballpark’ results that served as a good starting estimate for the model parameters. We also note that that while a formal inverse modeling approach would have been utilized, we found in pilot studies that manually adjusting the parameters to obtain a very good fit. This is because for the Yeoh model form the *C*_1_ term is dominant and adjusts the overall material stiffness, whereas changes to the *C*_2_ and *C*_3_ terms fine tune the nonlinear material behavior at higher strain levels. Thus, we specifically started by first estimating *C*_10_ from the analytical model response. Next, using the FE simulations, *C*_20_ and *C*_30_ underwent small adjustments until the best agreement between the model and experimental data was found. Using these calibrated parameters, we evaluated the predictive capabilities by simulating to predict the 1:3 and 3:1 biaxial extension mode responses.

#### RVE size effects

While we have rigorously established the optimal minimal RVE size, it is instructive to see how much an effect the use of smaller sizes actually have on the resulting simulations. This provides insights on how local structural variations can affect the resulting predictive stresses. This was done using the calibrated model parameters under a 1:1 biaxial extension for a 25 *µm*^3^, 50 *µm*^3^, and 75 *µm*^3^ cubic meshes. Simulation run times for the 25 *µm*^3^, 50 *µm*^3^, and 75 *µm*^3^ sized meshes completed within 30 minutes, 12 hours, and 16 hours, respectively.

### 2.8. Post-Processing

After completing all simulations, the effective strains and total reaction forces on each boundary face was determined for each load step. We then compared with the previous analytical model. Additionally, to compare the fiber mesh model of the previous analytical model in [12] with the finite element fiber mesh model, an additional volume mesh was generated in ABAQUS to represent a single electrospun fiber. The mesh was represented with a cylindrical geometry (10 *µ*m length, 1 *µ*m diameter) and C3D4 solid elements. Boundary conditions were chosen to simulate longitudinal extension of the fiber. The mesh material model was represented with the previously calibrated fiber material model parameters.

## 3. RESULTS

### 3.1. Simulation results

The final calibrated fiber mesh model for the 1:1 (equi-biaxial) deformation mode resulted in the following Yeoh material model parameters: *C*_1_ = 300 kPa, *C*_2_ = 100 kPa, *C*_3_ = 100 Pa, *D*_1_ = 41.6 kPa^−1^, *D*_2_ = 125 kPa^−1^, *D*_3_ = 125 kPa^−1^. Overall, the simulation results agreed well with the equi-biaxial experimental data (Figure 5), with *r*^2^ = 0.913 for *P*_11_ and r^2^ = 0.974 for *P*_22_. Using these same parameters, the stress responses were predicted for the 1:3 loading path, which showed overall good agreement with *P*_11_ r^2^ = 0.896 and R^2^ = 0.891 for *P*_22_ (Figure 6). For the 3:1 deformation model good agreement was also found, with r^2^ = 0.844 for *P*_11_ and r^2^ = 0.983 for *P*_22_ (Figure 7).

**Figure 5.**
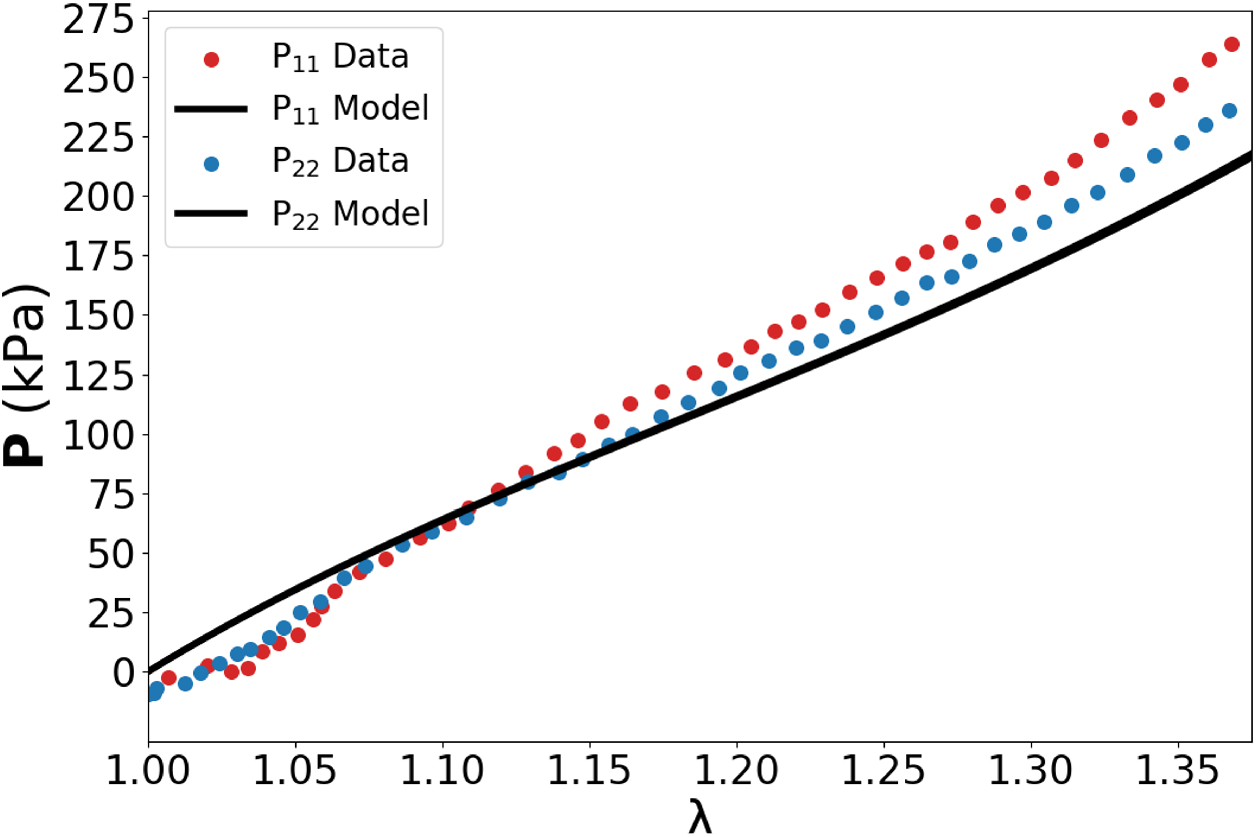
Stress v. stretch plot for biaxial 1:1 deformation mode using calibrated fiber mesh material model parameters.

**Figure 6.**
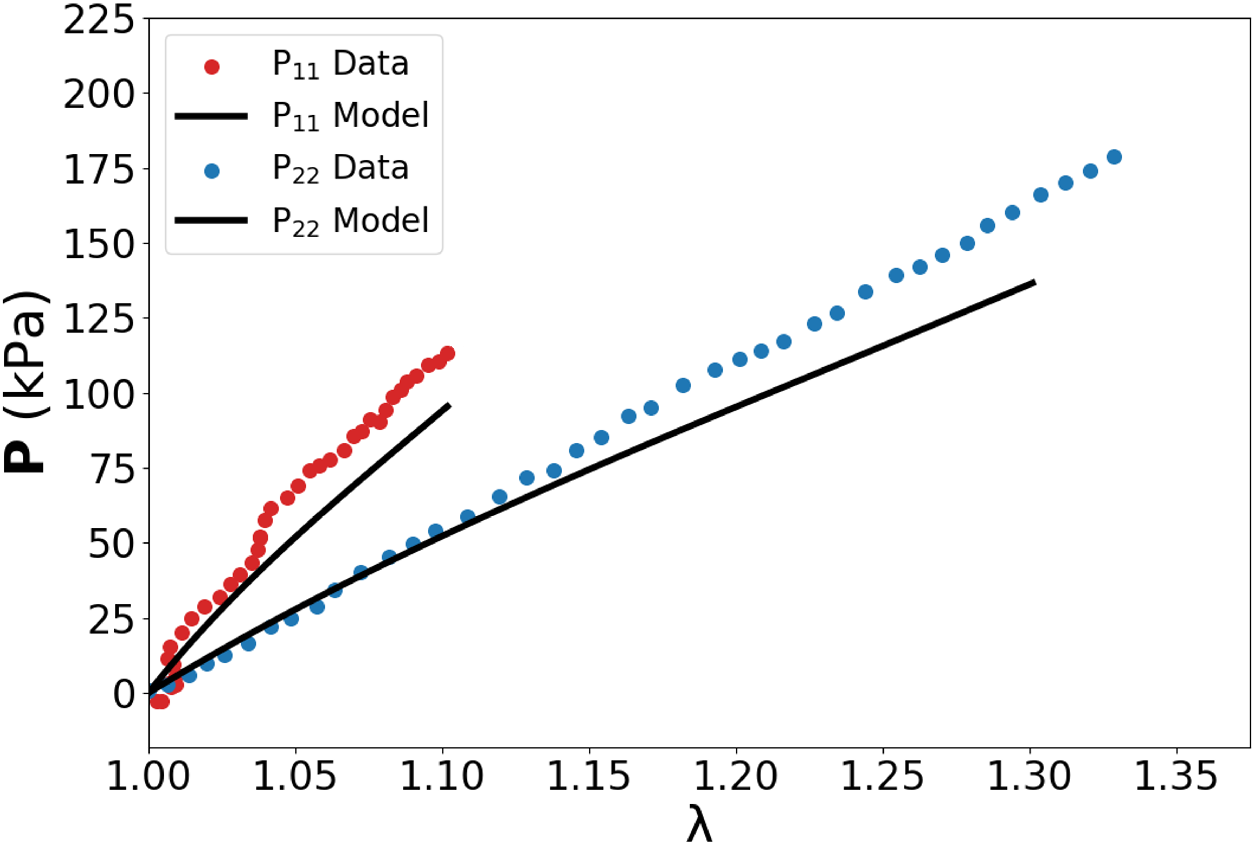
Stress v. stretch plot for biaxial 1:3 deformation mode using calibrated fiber mesh material model parameters.

**Figure 7.**
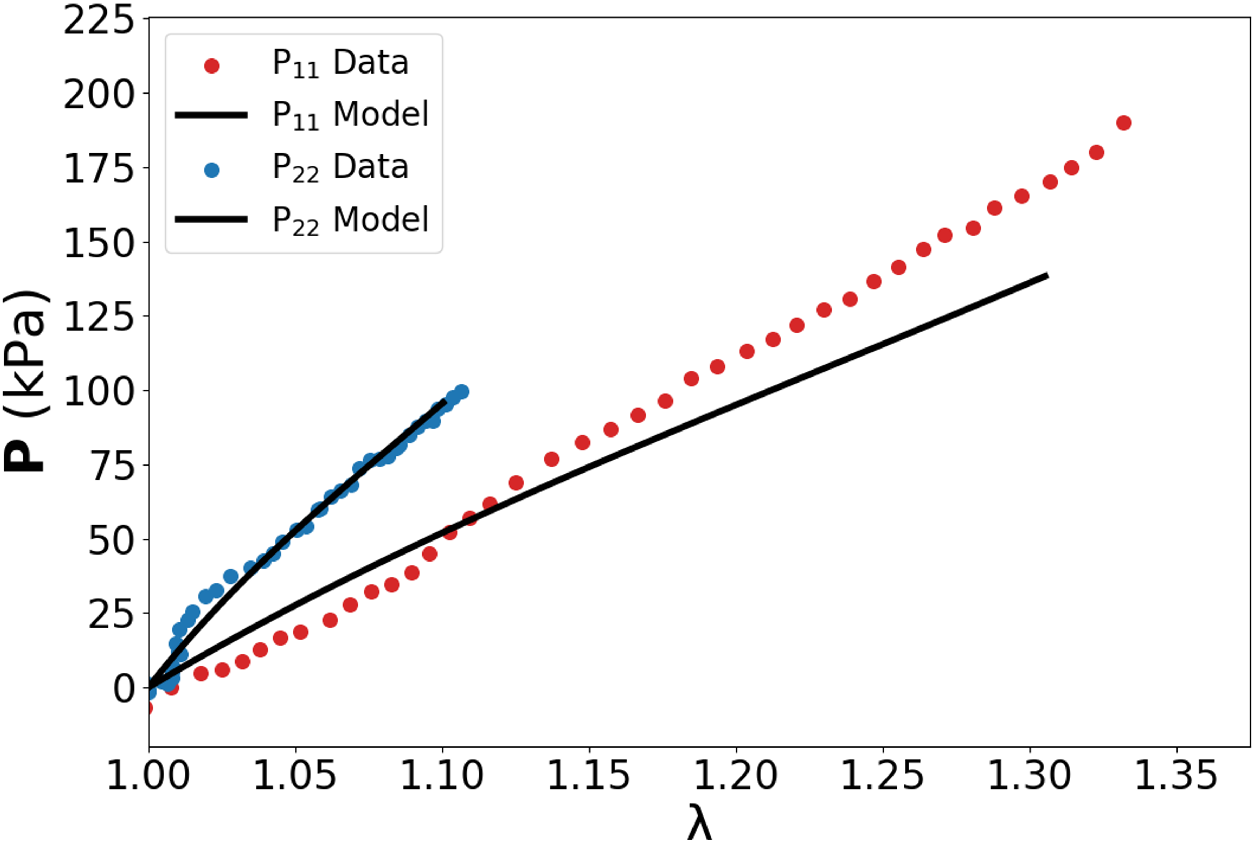
Stress v. stretch plot for biaxial 3:1 deformation mode using calibrated fiber mesh material model parameters.

### 3.2. FE simulation based vs analytical single fiber responses

A comparison between the finite element model and analytical model for a single fiber displaced uniaxially shows that that the finite element model (peak stress of 3.095 MPa) underpredicts the mechanical response, compared to the analytical model (peak stress of 5.853 MPa). The analytical model’s stress prediction also rises at a quicker rate than the stress prediction of the finite element model for larger extensions (Figure 8).

**Figure 8.**
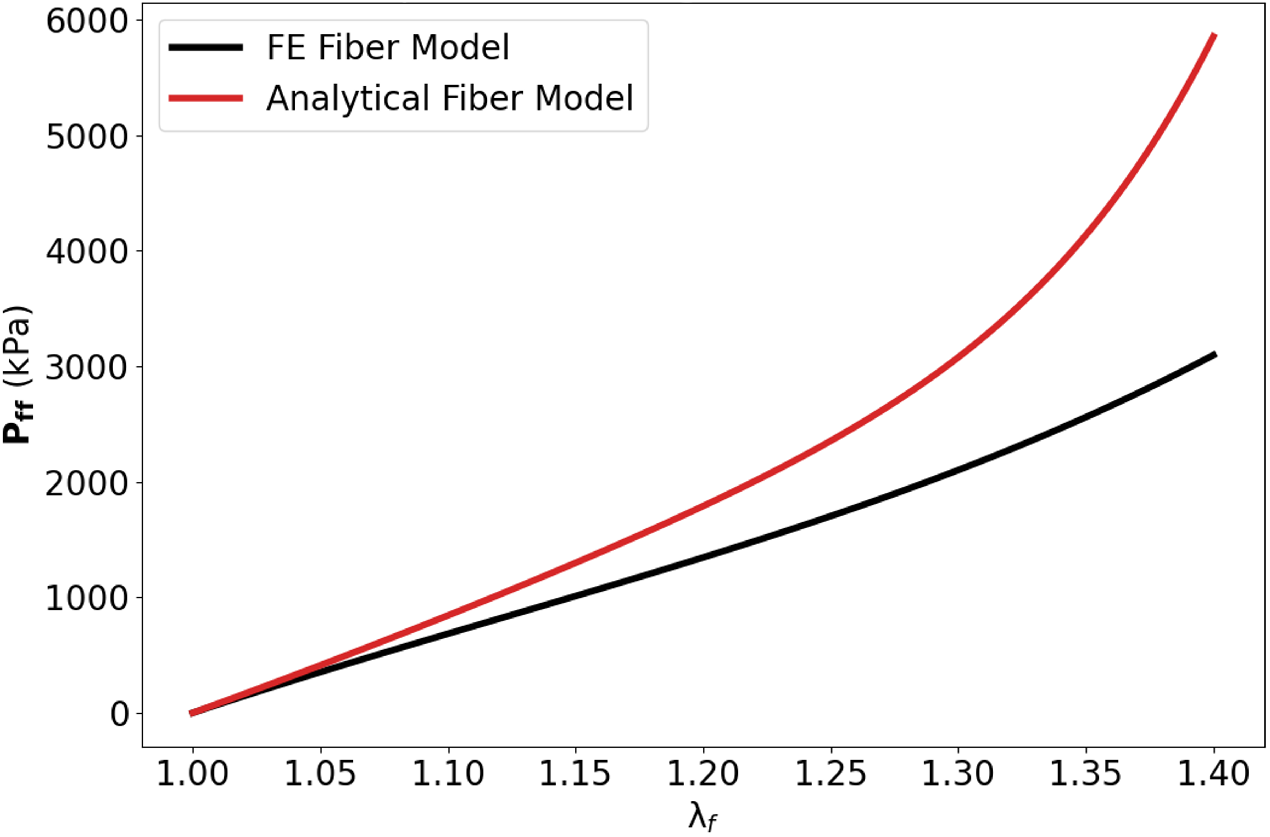
Uniaxial extension comparison between analytical fiber model and finite element model for a single fiber.

### 3.3. RVE Size effects

Comparing the 25 *µm*^3^, 50 *µm*^3^, and 75 *µm*^3^ cubic meshes for biaxial 1:1 deformation with experimental data shows that the 75 *µm*^3^ cubic mesh results consistently outperforms the smaller mesh sizes. The tradeoff for this accuracy was increased computational time; the 25 *µm*^3^ mesh taking 30 minutes, the 50 *µm*^3^ mesh taking 12 hours, and the 75 *µm*^3^ mesh taking 16 hours to complete (Figure 9).

**Figure 9.**
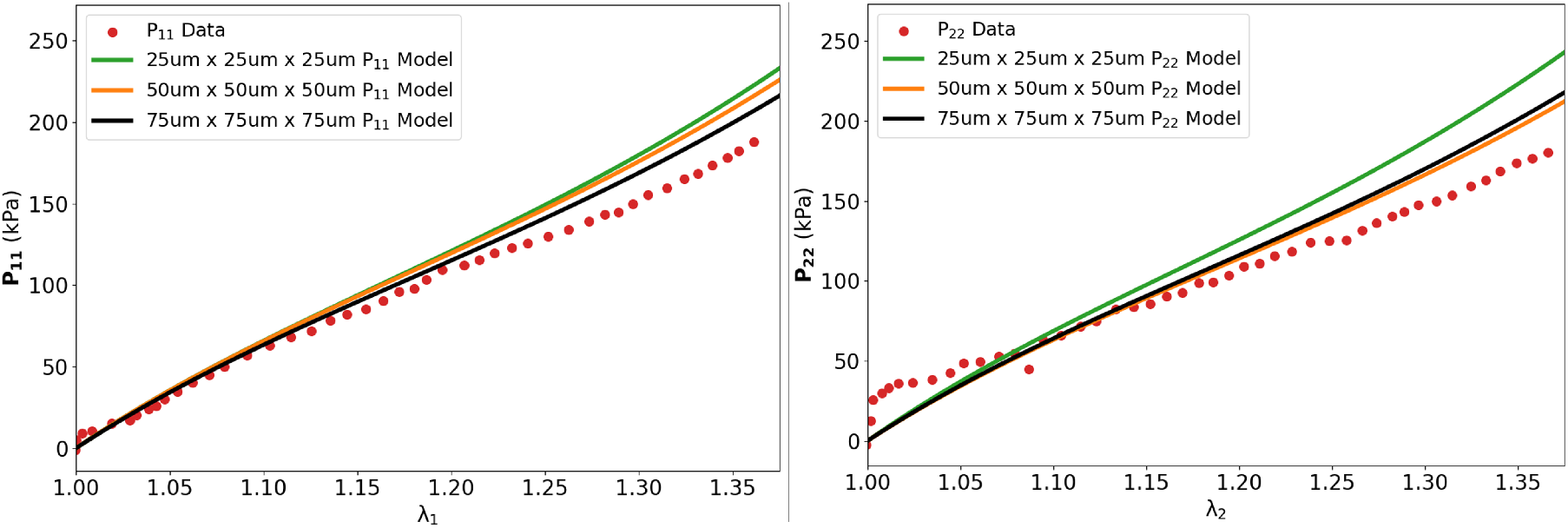
Biaxial 1:1 extensional deformation results for 25 *µm*^3^, 50 *µm*^3^, and 75 *µm*^3^ cubic meshes plot against experimental data. The 75 *µm*^3^ cubic mesh results are consistently closer to the experimental data measurements.

### 3.4. Local mechanical behaviors

In addition to reproducing stress-strain responses, the FE modeling approach allows visualizing the local mechanical behaviors of the fiber-hydrogel composite mesh. Inspecting these responses allows us to gain insight into how local fiber-fiber interactions may have contributed to the robust bulk mechanical properties, as found in [12]. Across the the biaxial 1:1, 1:3, and 3:1 deformation modes, all elements consistently showed that the maximum principal Cauchy stress in the fibers was roughly ten times larger than the maximum principal stress in hydrogel (Figure 10). This indicates that the fibers are the primary load-bearing component within the composite mesh model. In contrast, the maximum principal logarithmic strain for all elements in these simulations showed that the maximum principal strain in fiber is lower than the maximum principal strain in hydrogel (Figure 11). This indicates that the hydrogel undergoes larger deformations to accommodate the overall stretch, compared to the stiffer fibers. These findings demonstrate that the finite element model inherently captures the microscale behavior of the composite without the need for an additional interaction term. The material model definition and geometry of the fiber and hydrogel regions alone can accurately predict the bulk mechanical properties of the composite mesh. The higher observed principal stresses for fiber elements and higher observed principal strains for hydrogel elements accurately represent the stiffer fiber material model definition, compared to the more compliant hydrogel material model definition.

**Figure 10.**
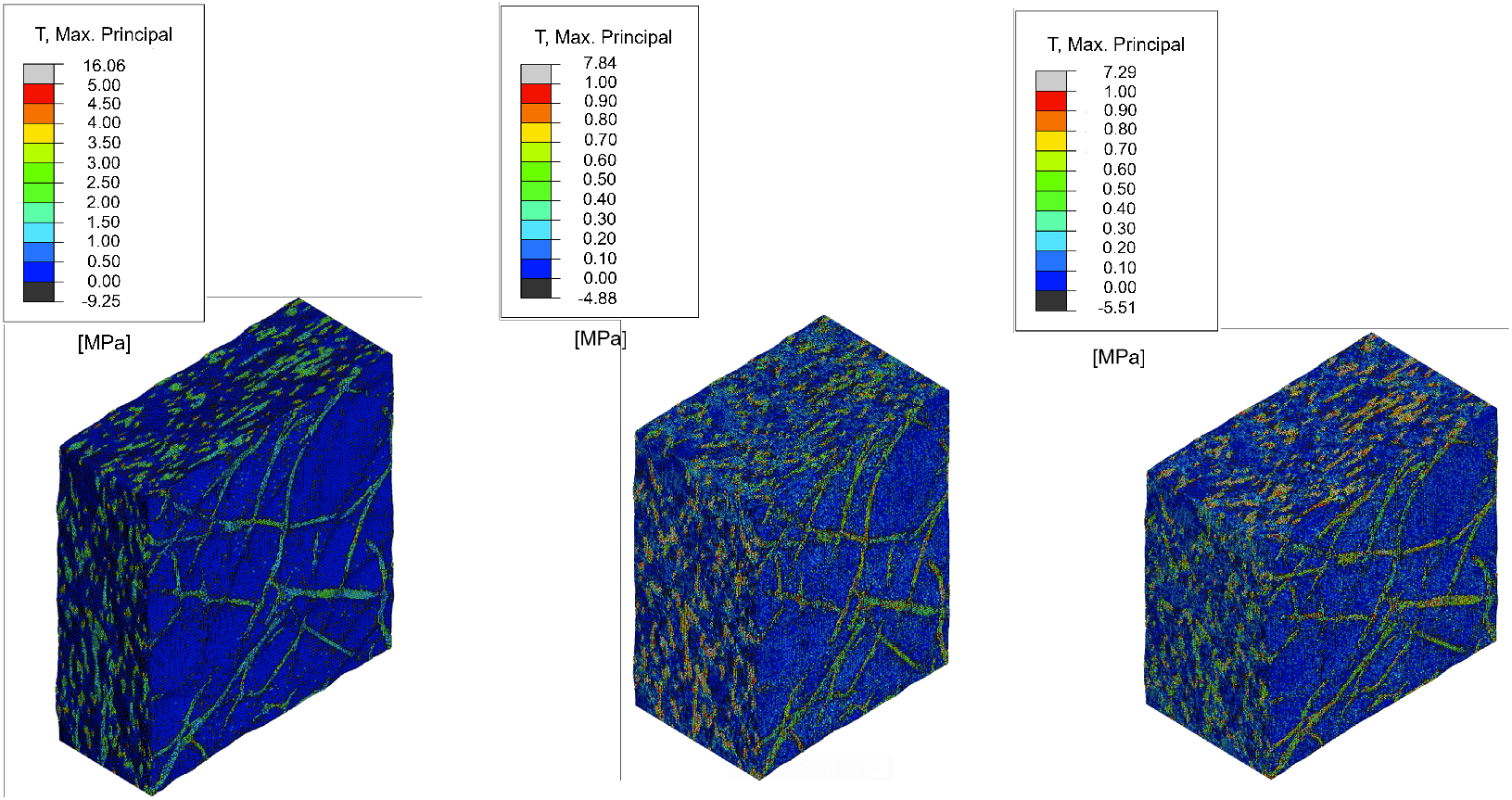
Maximum principal Cauchy stress distribution across biaxial 1:1 (left), biaxial 1:3 (center), and biaxial 3:1 (right) deformation simulations.

**Figure 11.**
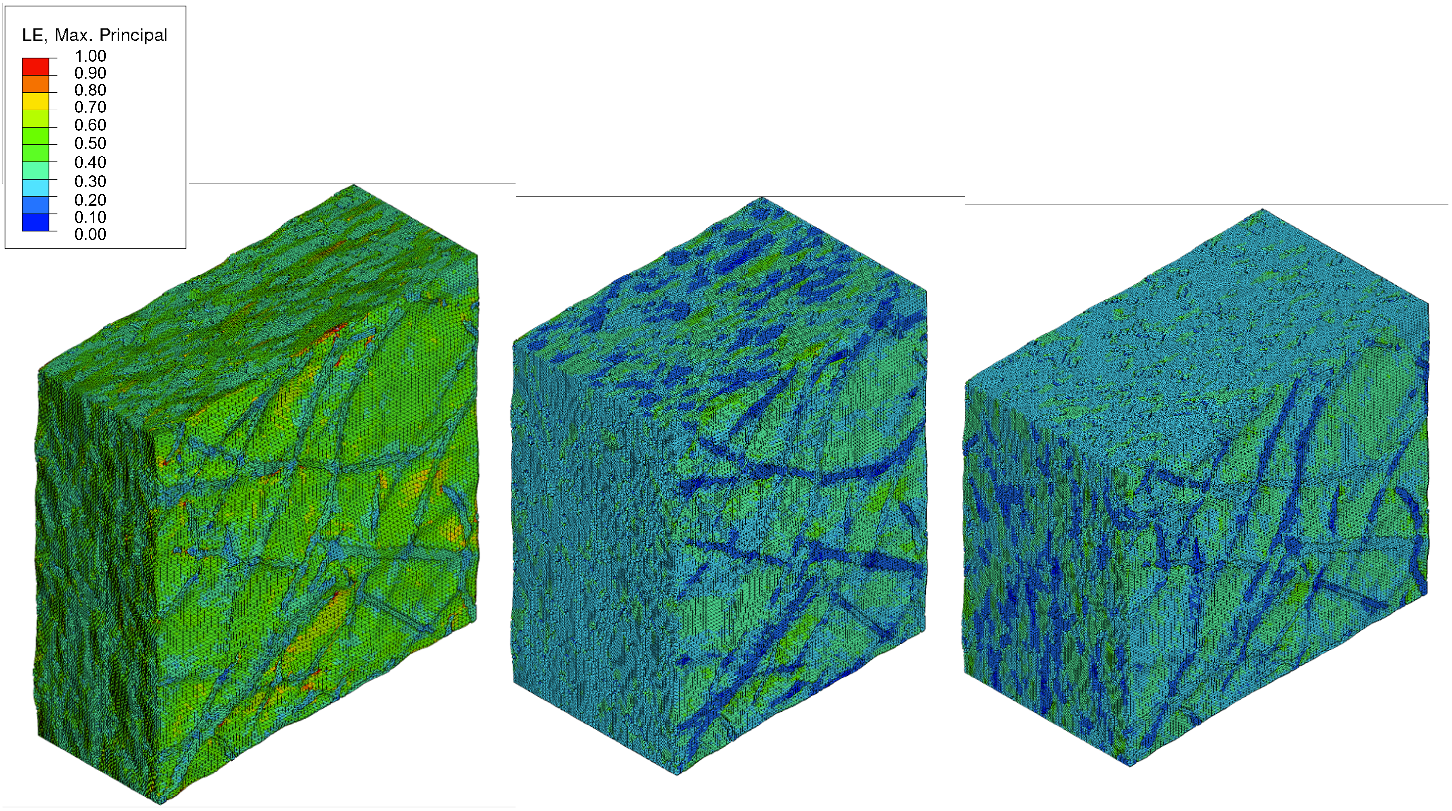
Maximum principal logarithmic strain distribution across biaxial 1:1 (left), biaxial 1:3 (center), and biaxial 3:1 (right) deformation simulations.

## 4. DISCUSSION

### 4.1. Key findings

In the present work, we developed a full 3D micromechanical model of electrospun meshes under biaxial loading to directly generate bulk mechanical behaviors. Our goal was to understand actual 3D fiber geometry at the microscale level impacts the bulk mechanical properties of fibrous electrospun meshes. The resultant simulations were then compared to experimental data reported by [12]. The appropriate RVE size that was calculated was found to be 75 *µm*^3^. The final calibrated fiber mesh model parameters were chosen based on the Biaxial 1:1 model prediction agreement with experimental data. Good agreement was dependent on a high coefficient of determination (R^2^ *>* 0.90). Both the Biaxial 1:3 and Biaxial 3:1 model predictions using the calibrated fiber mesh model parameters performed well against their experimental data measurements. As expected, the 75 *µm*^3^ cubic mesh outperformed both the 25 *µm*^3^ and 50 *µm*^3^ cubic mesh when comparing the results to experimental data. This further reinforces the RVE convergence analysis for finding an appropriate RVE size. Since the 25 *µm*^3^ and 50 *µm*^3^ cubic mesh sizes were consistently less accurate than the 75 *µm*^3^ cubic mesh, it can be concluded that a mesh size smaller than the appropriate RVE size cannot accurately represent the electrospun fiber mesh mechanical behavior as a whole.

In the present work, the use of actual 3D microgeometry allowed a more detail study, as well as incorporating a gel phase. The FE micro-model should thus simulate all fiber-gel interactions accurately, as well as fiber intersections in full 3D. It is interesting to note that the fiber-hydrogel composite material experiences the most stress within the fiber and the most strain within the hydrogel. This key result underscores that while the analytical model assumes local affine deformations [23], at the microscale this assumption does not hold. This result extends the results of Carleton et al. [7], who also found non-affine local deformations in planar 2D simulations using synthetic mesh geometries.

The complexity of the local responses (Figures 10,11) underscore the need for such approaches as undertaken here. Moreover, these local responses also manifest themselves at the bulk level. For example, while both the analytical and current FE models fit the experimental data well, their respective fiber responses were quite different (Figure 8), with the analytical model showing greater non-linearity and higher stresses overall. Given the high fidelity of the present approach, it is likely these differences are due to the substantial heterogeneous non-affine local deformations present in the actual electrospun fiber-hydrogel composite. Interestingly, no interaction term(s) were required by the FE model, and thus presents a more robust approach for these predictions. Since the finite element model is created using nanoCT images of the direct fiber geometry, this model is able to fully capture any previously assumed interactions occurring locally within the fiber mesh microstructure. This eliminated the need for adding any additional interaction mechanisms in the finite element model. Collectively, these findings further reveal the need for more realistic approaches if we are to better understand the how electrospun meshes actually function into order to improve their designs.

### 4.2. Limitations

While the current modeling approach predicted the experiments responses well overall, it did tend to under-predict the bulk mechanical response somewhat for the 1:3 and 3:1 protocols (Figures 6,7). This is most likely due to imperfections in the experiential setup, and naturally occurring variations in the local structure that were not captured in the current model. It is also possible that additional secondary micromechanical mechanisms exist, such as fiber bending/twisting not captured in the analytical model. Next, while our approach was able to glean such insights, the model imaging and development and the resulting simulations using ABAQUS were both time consuming. This limits the number of simulations that can be developed and run in a reasonable time frame. However, in recent years there have significant development in automated segmentation and high speed computational methods for hyperelastic and biomechanical applications (based on advanced machine-learning based approaches) [24, 25], will make these approaches much more practical.

### 4.3. Potential Applications

The present study was aimed at improving our understanding of these fiber-hydrogel composites at the microscale. One very relevant application of gel embedded ES meshes are replacement heart valves (RHV) biomaterials. Current RHV xenograft biomaterials, even with anticalcification treatments [26–28], remain problematic to address the needs for other adult heart valves and the pediatric population. A new generation of heart valve biomaterials is clearly needed that are i) durable, ii) hemocompatible, iii) allow sufficient variations in fabrication to produce a wide range of mechanical responses, and iv) allow for superior hemodynamics in a device setting. This is a substantial challenge; material properties which promote the requisite hemocompatibility may not be consistent with those required for durability or hemodynamics. To decouple these design goals, electrospun fiber-hydrogel composites show promise, as the surface hydrogel provides thromboresistance and bacterial resistance, while an electrospun fiber mesh provides the requisite mechanical properties.

Clearly, hierarchical control of the polymer structure from the molecular scale to the microarchitectural features of the fibrous mesh potentially provides a highly tunable design space to achieve target properties for a wide range of RHV leaflet designs. As these responses are governed by underlying fiber orientation dictated by the fabrication process, they can be designed. Interestingly, both the previous a current study we also observed no evidence of fiber-hydrogel mechanical interactions. This novel finding suggests that the constituent electrospun mesh microarchitectures and hydrogel blood contacting properties can be optimized *independently*, allowing for a much wider range of design possibilities. The present approach can be extended to include multi-scale computational methods

### 4.4. Conclusions

Our goal herein was to understand how actual 3D fiber geometry at the microscale level impacts the bulk mechanical properties of electrospun fiber-hydrogel composites. Our framework enabled systematic investigation of both the macroscopic bulk mechanical response of the overall fiber mesh and the microscopic localized mechanical response of fibers under various stages of loading. The resultant simulations were generally accurate and predictive. RVE size studies indicated an 75 *µ*m length scale for simulations, which was corroborated by simulations run at smaller sizes that revealed greater variations. Interestingly, in addition matching the experimental bulk stress-strain responses, we also found very different effective fiber stress-strain responses from the previous analytical model. It is likely these differences are due to the substantial heterogeneous non-affine local deformations observed in the simulations of the actual electrospun fiber-hydrogel composites. This finding further revealed the need for more rigorous approaches to better understand how electrospun meshes function in order to improve their use in modern medical devices and implants.

## 6. Acknowledgments

The authors gratefully acknowledge support of NIH/NHLBI grant R01HL175284 to MSS and ECH. Bionate® 80A was provided by the biomedical division of dsm-firmenich (Exton, PA).

## References

[1] N. Bhardwaj, S. C. Kundu, Electrospinning: A fascinating fiber fabrication technique, Biotechnology Advances 28 (3) (2010) 325–347. doi:10.1016/j.biotechadv.2010.01.004. URL https://www.sciencedirect.com/science/article/pii/S0734975010000066

[2] S. Heydarkhan-Hagvall, K. Schenke-Layland, A. P. Dhanasopon, F. Rofail, H. Smith, B. M. Wu, R. Shemin, R. E. Beygui, W. R. MacLellan, Three-dimensional electrospun ecm-based hybrid scaffolds for cardiovascular tissue engineering, Biomaterials 29 (19) (2008) 2907–2914. doi:10.1016/j.biomaterials.2008.03.034. URL https://www.sciencedirect.com/science/article/pii/S0142961208002068

[3] P.-h. Grace Chao, H.-Y. Hsu, H.-Y. Tseng, Electrospun microcrimped fibers with nonlinear mechanical properties enhance ligament fibroblast phenotype, Biofabrication 6 (3) (2014) 035008. doi:10.1088/1758-5082/6/3/035008. URL 10.1088/1758-5082/6/3/035008

[4] T. Courtney, M. S. Sacks, J. Stankus, J. Guan, W. R. Wagner, Design and analysis of tissue engineering scaffolds that mimic soft tissue mechanical anisotropy, Biomaterials 27 (19) (2006) 3631–3638. doi:10.1016/j.biomaterials.2006.02.024. URL https://www.sciencedirect.com/science/article/pii/S0142961206001645

[5] A. Stylianopoulos, C. Bashur, A. Goldstein, S. Guelcher, V. Barocas, Computational predictions of the tensile properties of electrospun fibre meshes: Effect of fibre diameter and fibre orientation, Journal of Mechanical Behavior of Biomedical Materials 1 (2008) 326–335.

[6] D’Amore J. A. Stella, W. R. Wagner, M. S. Sacks, Characterization of the complete fiber network topology of planar fibrous tissues and scaffolds, Biomaterials 31 (20) (2010) 5345–54. doi:10.1016/j.biomaterials.2010.03.052. URL http://www.ncbi.nlm.nih.gov/entrez/query.fcgi?cmd=Retrieve&db=PubMed&dopt=Citation&list_uids=20398930

[7] J. B. Carleton, A. D’Amore, K. R. Feaver, G. J. Rodin, M. S. Sacks, Geometric characterization and simulation of planar layered elastomeric fibrous biomaterials., Acta biomaterialia 12 (2015) 93–101. doi:10.1016/j.actbio.2014.09.049.

[8] J. B. Carleton, A. D’Amore, K. R. Feaver, G. J. Rodin, M. S. Sacks, Geometric characterization and simulation of planar layered elastomeric fibrous biomaterials, Acta Biomaterialia 12 (10 2014). doi:10.1016/j.actbio.2014.09.049.

[9] M. Rizvi, P. Kumar, D. Katti, A. Pal, Mathematical model of mechanical behavior of micro/nanofibrous materials designed for extracellular matrix substitutes, Acta Biomaterialia 8 (11) (2012) 4111–4122. doi:10.1016/j.actbio.2012.07.025. URL https://www.sciencedirect.com/science/article/pii/S1742706112003443

[10] W. Zhang, S. Motiwale, M.-C. Hsu, M. S. Sacks, Simulating the time evolving geometry, mechanical properties, and fibrous structure of bioprosthetic heart valve leaflets under cyclic loading. 123 104745. doi:10.1016/j.jmbbm.2021.104745.

[11] S. Motiwale, M. D. Russell, O. Conroy, J. Carruth, M. Wancura, A. Robinson, E. Cosgriff-Hernandez, M. S. Sacks, Anisotropic elastic behavior of a hydrogel-coated electrospun polyurethane: Suitability for heart valve leaflets. 125 104877. doi:10.1016/j.jmbbm.2021.104877.

[12] S. Motiwale, M. D. Russell, O. Conroy, J. Carruth, M. Wancura, A. Robinson, E. Cosgriff-Hernandez, M. S. Sacks, Anisotropic elastic behavior of a hydrogel-coated electrospun polyurethane: Suitability for heart valve leaflets, Journal of the mechanical behavior of biomedical materials 125 (2022) 104877.

[13] M. S. Sacks, W. Zhang, S. Wognum, A novel fibre-ensemble level constitutive model for exogenous cross-linked collagenous tissues, Interface Focus 6 (1) (2016) 20150090. doi:10.1098/rsfs.2015.0090. URL http://rsfs.royalsocietypublishing.org/royfocus/6/1/20150090.full.pdf

[14] A. Schneider, W. S. Rasband, K. W. Eliceiri, NIH image to ImageJ: 25 years of image analysis, Nat. Methods 9 (7) (2012) 671–675.

[15] M. Hori, S. Nemat-Nasser, On two micromechanics theories for determining micro–macro relations in heterogeneous solids, Mechanics of Materials 31 (10) (1999) 667–682. doi:10.1016/S0167-6636(99)00020-4. URL https://www.sciencedirect.com/science/article/pii/S0167663699000204

[16] P. A. Wijeratne, V. Vavourakis, J. H. Hipwell, C. Voutouri, P. Papageorgis, T. Stylianopoulos, A. Evans, D. J. Hawkes, Multiscale modelling of solid tumour growth: The effect of collagen micromechanics, Biomechanics and modeling in mechanobiology (Oct 2016). URL https://pmc.ncbi.nlm.nih.gov/articles/PMC4762195/#R21

[17] J. Aboudi, S. M. Arnold, B. A. Bednarcyk, Micromechanics of composite materials: a generalized multiscale analysis approach, Butterworth-Heinemann, 2012.

[18] S. Bargmann, B. Klusemann, J. Markmann, J. E. Schnabel, K. Schneider, C. Soyarslan, J. Wilmers, Generation of 3d representative volume elements for heterogeneous materials: A review, Progress in Materials Science 96 (2018) 322–384. doi:10.1016/j.pmatsci.2018.02.003. URL https://www.sciencedirect.com/science/article/pii/S0079642518300161

[19] Comet Technologies Canada Inc., Dragonfly 2024.1 [computer software], https://dragonfly.comet.tech/, software available at https://dragonfly.comet.tech/ (2024).

[20] Preim, C. Botha, Chapter 4 - Image Analysis for Medical Visualization, second edition Edition, Morgan Kaufmann, Boston, 2014. doi:10.1016/B978-0-12-415873-3.00004-3. URL https://www.sciencedirect.com/science/article/pii/B9780124158733000043

[21] Geuzaine, J. Remacle, Gmsh: A 3-D finite element mesh generator with built-in preand post-processing facilities, International Journal for Numerical Methods in Engineering 79 (11) (2009) 1309–1331. doi:10.1002/nme.2579.

[22] Dassault Systèmes, ABAQUS 6.12 Theory Manual, http://abaqus.software.polimi.it/v6.12/books/stm/default.htm

[23] R. Fan, M. S. Sacks, Simulation of planar soft tissues using a structural constitutive model: Finite element implementation and validation, Journal of Biomechanics 47 (2014) 2043–2054.

[24] S. Motiwale, W. Zhang, M. S. Sacks, High-Speed High-Fidelity Cardiac Simulations Using a Neural Network Finite Element Approach, in: O. Bernard, P. Clarysse, N. Duchateau, J. Ohayon, M. Viallon (Eds.), Functional Imaging and Modeling of the Heart, Springer Nature Switzerland, pp. 537–544. doi:10.1007/978-3-031-35302-4_55.

[25] M. S. Sacks, S. Motiwale, C. Goodbrake, W. Zhang, Neural network approaches for soft biological tissue and organ simulations, Journal of Biomechanical Engineering 144 (12) (2022) 121010.

[26] M. N. Girardot, M. Torrianni, J. M. Girardot, Effect of AOA on glutaraldehyde-fixed bioprosthetic heart vavlve cusps and walls: binding and calcification studies, Int J Artif Organs 17 (1994) 76–82.

[27] J. P. Gott, M. N. Girardot, J. M. Girardot, J. D. Hall, J. D. Whitlark, W. S. Horsley, L. M. Dorsey, R. J. Levy, W. Chen, F. J. Schoen, R. A. Guyton, Refinement of the alpha aminooleic acid bioprosthetic valve anticalcification technique, Ann Thorac Surg 64 (1) (1997) 50–8. URL http://www.ncbi.nlm.nih.gov/entrez/query.fcgi?cmd=Retrieve&db=PubMed&dopt=Citation&list_uids=9236334

[28] D.J. Myers, G. Nakaya, M. N. Girardot, G. W. Christie, A comparison between glutaraldehyde and diepoxide-fixed stentless porcine aortic valves: biochemical and mechanical characterization and resistance to mineralization, J Heart Valve Dis 4 Suppl 1 (1995) S98–101.

